# A tripartite mechanism licenses RNA polymerase import into the phage nucleus

**DOI:** 10.64898/2026.09.21.753271

**Authors:** Claire Kokontis, Deepto Mozumdar, Daphne Chen, Iris Zheng, Wearn-Xin Yee, David Bulkley, David A. Agard, Joseph Bondy-Denomy

## Abstract

Nucleus-forming jumbo phages import a phage-encoded non-virion RNA polymerase (nvRNAP) into a proteinaceous nucleus, but how it is selected for import is unknown. Here, we identify an import pathway that couples nvRNAP subunit interactions with phage-encoded import factors. Maximal nvRNAP import requires a novel factor Imp7 (gp166) and the essential import factor Imp1. Imp7 is essential at environmental temperatures (20 °C), where impaired nvRNAP import causes a profound loss of middle and late transcription and blocks phage DNA replication. Import of individual nvRNAP subunits also depends on other subunits, consistent with import licensing at the level of an assembled complex. In the absence of middle transcription and DNA replication, the phage nucleus surprisingly still assembles and segregates from the cytoplasm, demonstrating that nvRNAP nuclear localization and activity are not a critical checkpoint for nucleus assembly. Together, these findings define a specialized pathway for nuclear import of the multi-subunit RNAP and reveal that nuclear compartment assembly precedes the transcriptional and replicative programs required for its maturation.

## Introduction

The phage ΦKZ evades host nucleases by segregating its genome in a series of three selectively permeable compartments: a lipid vesicle, a proteinaceous phage nucleus, and ultimately the phage capsid. Upon infection initiation, the ΦKZ genome is injected into the lipid-bound early phage infection (EPI) vesicle with the phage-encoded multi-subunit virion RNA polymerase (vRNAP)^1–6^. The vRNAP transcribes early phage genes inside the EPI vesicle^1–3,7^, which are then translated outside the vesicle^3^. The phage genome is then transferred to the nascent proteinaceous phage nucleus composed primarily of ChmA (gp54)^8–10^. A second RNA polymerase, the non-virion RNAP (nvRNAP), is transcribed by the vRNAP early during infection^7^, translated in the cytoplasm, and imported into the phage nucleus through an unknown mechanism to execute middle/late transcription. vRNAP subunits are excluded from the nucleus and remain with the vesicle^1,2,8^. Genes encoding proteins involved in DNA replication (e.g., phage-encoded DNA polymerase, RecA/UvsX recombinase) are transcribed inside the phage nucleus^11^, translated in the cytoplasm, and are subsequently imported^8,12,13^. Phage nucleus maturation then occurs alongside middle transcription and subsequent phage DNA replication^8,9,11,14,15^.

The import of nuclear proteins, including both phage- and host-encoded DNA replication-associated proteins and a phage antagonist of defense system DarT2^16^, requires essential and conserved protein Imp1^13,17^. Imp1 localizes to the nascent nucleus early in infection and remains at the periphery of the mature nucleus throughout infection^13,17^. Imp1 uses cargo-specific interfaces to recognize and engage with nuclear cargo to facilitate their import into the phage nucleus^13^. Cargo-specific adaptors Imp2, Imp4, and Imp5 are required for the import of specific proteins^13^. It remains unknown whether the essential multi-subunit nvRNAP similarly requires Imp1 interfaces or any other factors to achieve specific import.

Here, we performed an unbiased genetic selection to identify the nuclear import requirements for the phage nvRNAP. We identified Imp7 (gp166) as a novel nvRNAP import factor, which is required for maximal nvRNAP import into the phage nucleus. nvRNAP subunit-subunit interactions also appear to be critical for import. The import of Imp7 itself also requires Imp1 and Imp3, previously identified essential import factors^13,18^. Interestingly, Imp7 is conditionally essential at environmental temperatures (18-20 °C) where it is required for Imp1-dependent nvRNAP import. When the phage is starved for nvRNAP import, phage DNA fails to replicate, while the phage nucleus surprisingly still partially assembles, segregates phage nuclear contents from the cytoplasm and is centered by PhuZ. These data reveal that import of the essential multi-subunit nvRNAP is achieved through participation of at least three different classes of phage proteins, but internalization and activity of this complex is not required for nuclear shell assembly.

## Results

### nvRNAP import requires Imp7 and subunit-subunit interactions

The nvRNAP is composed of gp74 and a spliced fusion of gp55 and gp56.1 (together comprising the split β’ subunits, hereafter referred to as gp55), gp123 and a spliced fusion of gp71 and gp73 (the split β subunits), and gp68 (σ factor)^7,19–21^. Activity of nvRNAP is essential according to phage transposon mutagenesis^18^, ASO knockdown^11^, and our isolation of a temperature sensitive mutant (*ts1*) with an *E244K* mutation in *orf74*, which fails to replicate at the nonpermissive temperature (41 °C) (Sup Fig 1a). To identify the genes required for import of the nvRNAP proteins, we selected for phage mutants that decrease the import of EcoRI-sfCherry2 fusions to each of the five nvRNAP subunits (hereafter referred to as EcoRI-nvRNAP fusions). EcoRI fusions to gp55, gp68, or gp74 reduced ΦKZ titer by ∼10^4^-10^7^-fold (Fig 1a, Sup Fig 1b), revealing rare escape phage mutants that likely decrease import of nvRNAP subunits. EcoRI fusions to gp71-73 or gp123 were not active against WT ΦKZ (Sup Fig 1c).

**Figure 1:**
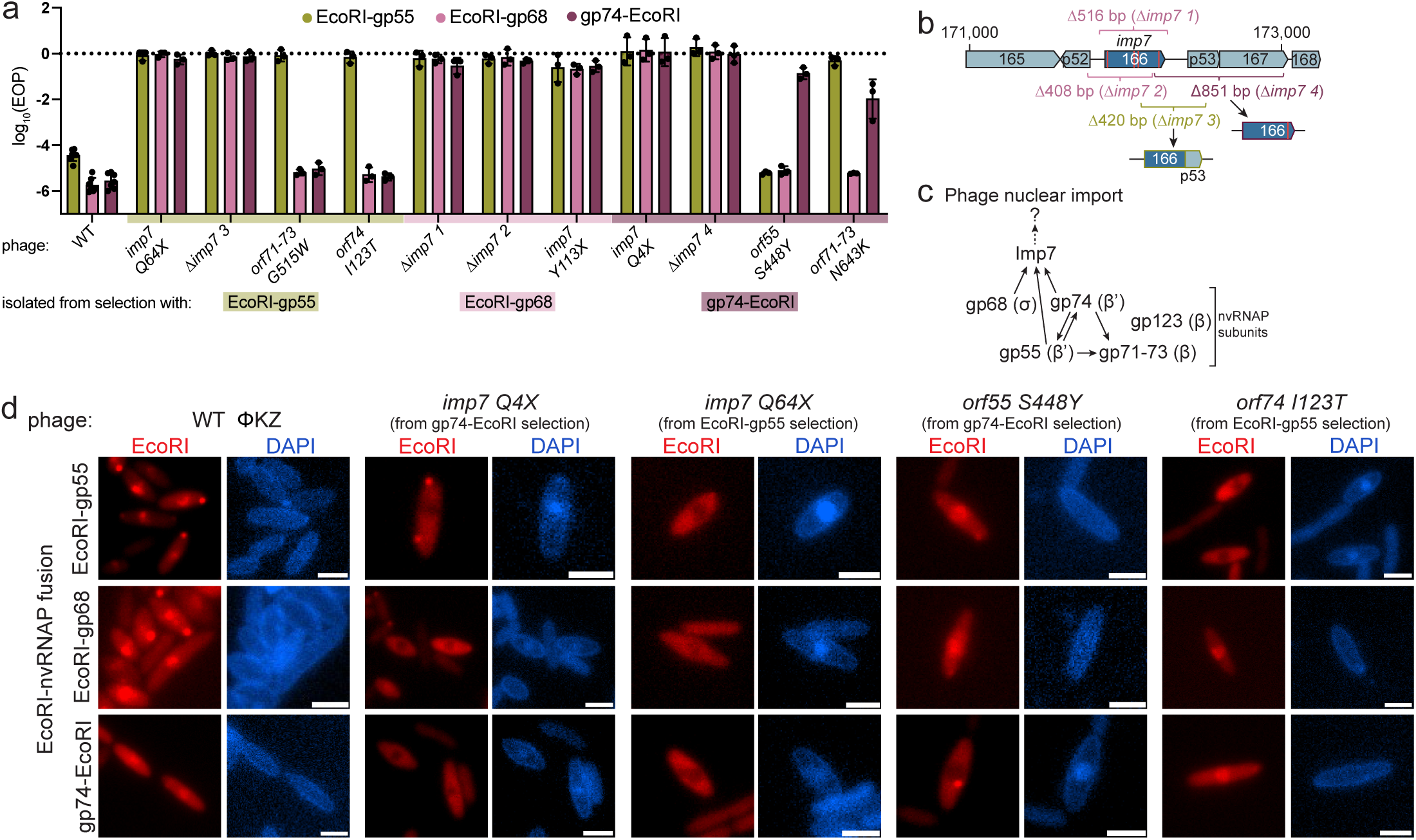
Mutations in *imp7* and nvRNAP subunits reduce import of nvRNAP subunits. a, log_10_ efficiency of plating (EOP) of indicated wildtype and EcoRI-nvRNAP escaper phages on EcoRI-nvRNAP expressing PAO1 strains. Plaquing efficiency is normalized to phage titer on a strain expressing non-targeting catalytically dead EcoRI-gp68. x-axis is colored to indicate which EcoRI-nvRNAP selection the phage was isolated from. b, Graphical description of *imp7* nonsense mutations and deletions isolated from selection. Red lines through *imp7* represent nonsense mutations. Colors of deletion labels indicate which EcoRI-nvRNAP selection the deletion was isolated from. c, Schematic representing the factors required for nuclear import of proteins queried in EcoRI selections, as determined by the escape mutations isolated in *imp7* and nvRNAP subunits from each selection (i.e. gp68 → Imp7 indicates gp68 requires wildtype Imp7 for its nuclear import). d, Live-cell fluorescence microscopy of wildtype or indicated mutant phage infecting PAO1 strain expressing the indicated EcoRI-sfCherry2-nvRNAP (referred to as EcoRI-nvRNAP) subunit fusion (red). DAPI (blue) indicates phage DNA. Scale bars represent 2 μm. WT ΦKZ: EcoRI-gp55 n=92/93 imported, EcoRI-gp68 n=91/91 imported, gp74-EcoRI n=97/97 imported. *imp7 Q4X*: EcoRI-gp55 n=90/95 reduced import, EcoRI-gp68 n=224/237 reduced import, gp74-EcoRI n=134/137 reduced import. *imp7 Q64X*: EcoRI-gp55 n=67/68 reduced import, EcoRI-gp68 n=138/150 reduced import, gp74-EcoRI n=77/79 reduced import. *orf55 S448Y*: EcoRI-gp55 n=92/92 imported, EcoRI-gp68 n=199/201 imported, gp74-EcoRI n=146/174 reduced import. *orf74 I123T*: EcoRI-gp55 n=110/128 reduced import, EcoRI-gp68 n=593/600 imported, gp74-EcoRI n=188/189 imported. Plaque assays were performed at least three independent times in biological replicates with similar results. Microscopy was performed at least two independent times in biological replicates with similar results.

Two distinct classes of phage mutants emerged that resisted the EcoRI fusion constructs. The first class consisted of escape phages with spontaneous mutations in *orf166,* an uncharacterized gene that we named *imp7* for import factor 7. *imp7* mutations encompassed nonsense and missense point mutations and deletions of *imp7* (Fig 1b, Table 1, Sup Table 1; unique imp7 deletions and truncations are hereafter referred to as Δ*imp7 1-4*). Imp7 is a small 15 kDa protein, and the AlphaFold 3 predicted structure of Imp7 is low confidence (pTM = 0.35) (Sup Fig 1d). By amino acid sequence, Imp7 is well-conserved only in *Pseudomonas*-infecting jumbo phages closely related to ΦKZ and has no sequence or structural homologs with known function.

**Table 1:**
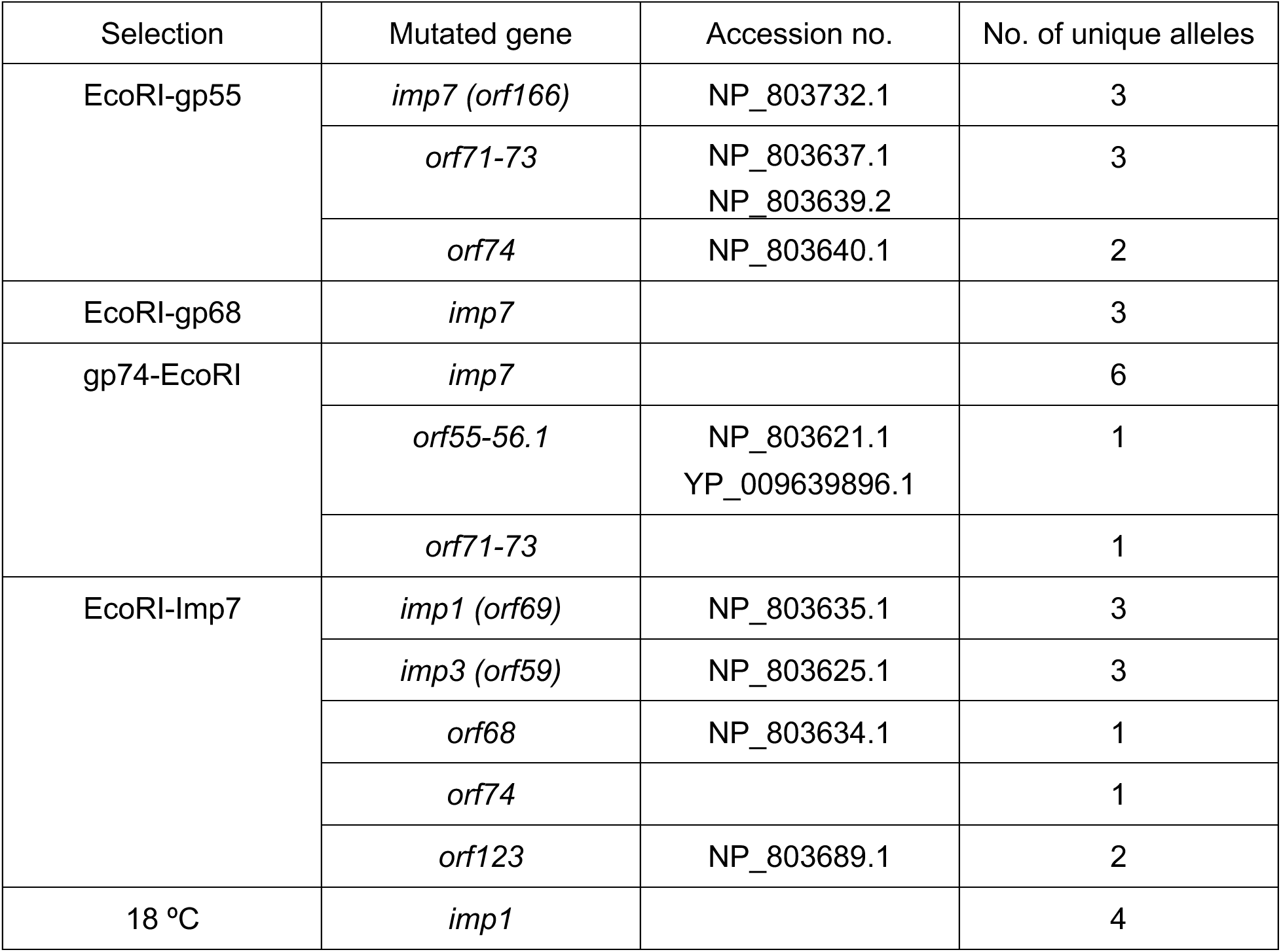
Summary of mutant alleles isolated from genetic selections. Summary of ΦKZ genes mutated under selection by the indicated EcoRI fusion or restrictive temperature condition, including gene accession numbers and how many unique mutations were isolated.

*imp7* mutant phages resist targeting by all three EcoRI-nvRNAP subunit fusions (Fig 1a). A clean *imp7::acrVIA1* replacement (hereafter referred to as *Δimp7*) in a WT ΦKZ background also resisted all three EcoRI-nvRNAP subunit fusions (Sup Fig 1e). Expression of Imp7 *in trans* resensitized *imp7* mutant phages to EcoRI-nvRNAP targeting, confirming *imp7* mutations are causal for decreased EcoRI-nvRNAP sensitivity (Sup Fig 1f). Live-cell fluorescence microscopy of wild type and *imp7* mutant infections confirmed that *imp7* mutants reduced import of all three EcoRI-nvRNAP fusions into the phage nucleus (Fig 1d), regardless of the EcoRI selection the mutant was isolated under. Together, these data demonstrate that *imp7* is required for maximal import of multiple nvRNAP subunits. This suggests that perhaps nvRNAP subunits are being imported or localized into the nucleus as a complex and thus have a single co-dependency.

The second class of mutants consisted of phages that acquired a single missense mutation in one of the subunits of the nvRNAP complex (Fig 1a, c, Sup Fig 1g, Table 1). For example, the EcoRI-gp55 fusion selected for missense point mutations in gp71-73 or gp74 (Fig 1a, c). As another example, the gp74-EcoRI fusion selected for missense point mutations in gp55 or gp71-73 (Fig 1a). In contrast to *imp7* mutations, nvRNAP subunit mutations conferred resistance to the EcoRI fusion they were selected under and maintained sensitivity to the two other EcoRI fusions (Fig 1a). For example, *orf55 S448Y* conferred resistance to only gp74-EcoRI, while *orf71-73 G515W* conferred resistance to only EcoRI-gp55. One exception was an *orf71-73 N643K* mutant selected by gp74-EcoRI that was resistant to both gp74-EcoRI and EcoRI-gp55, but not EcoRI-gp68, suggesting this particular gp71-73 residue is important for import of both gp55 and gp74 (Fig 1a). Expression of relevant wildtype nvRNAP subunits *in trans* resensitized mutant phages to EcoRI-nvRNAP targeting, confirming causality of these mutations for decreased EcoRI-nvRNAP sensitivity (Sup Fig 1f). We confirmed the subunit-specific import defects with live-cell fluorescence microscopy (Fig 1d). These results demonstrate that import of individual nvRNAP subunits can depend on other intact subunits, consistent with recognition and/or import of an assembled or partially assembled nvRNAP complex.

Next, we asked whether Imp7 mediates import of the nvRNAP by binding directly to the complex. We combined purified 6xHis-tagged Imp7 with 6xHis-tagged five subunit nvRNAP or 6xHis-tagged four subunit complex lacking gp68, which is stable *in vitro*^20^. We ran the mixtures over size-exclusion chromatography to assess binding and found that Imp7 did not stably bind to either the four subunit (without gp68) or five subunit complexes (Sup Fig 2a). These data suggest that although Imp7 is required for nvRNAP import, it executes this function through a non-direct interaction or that other proteins are required for binding.

**Figure 2:**
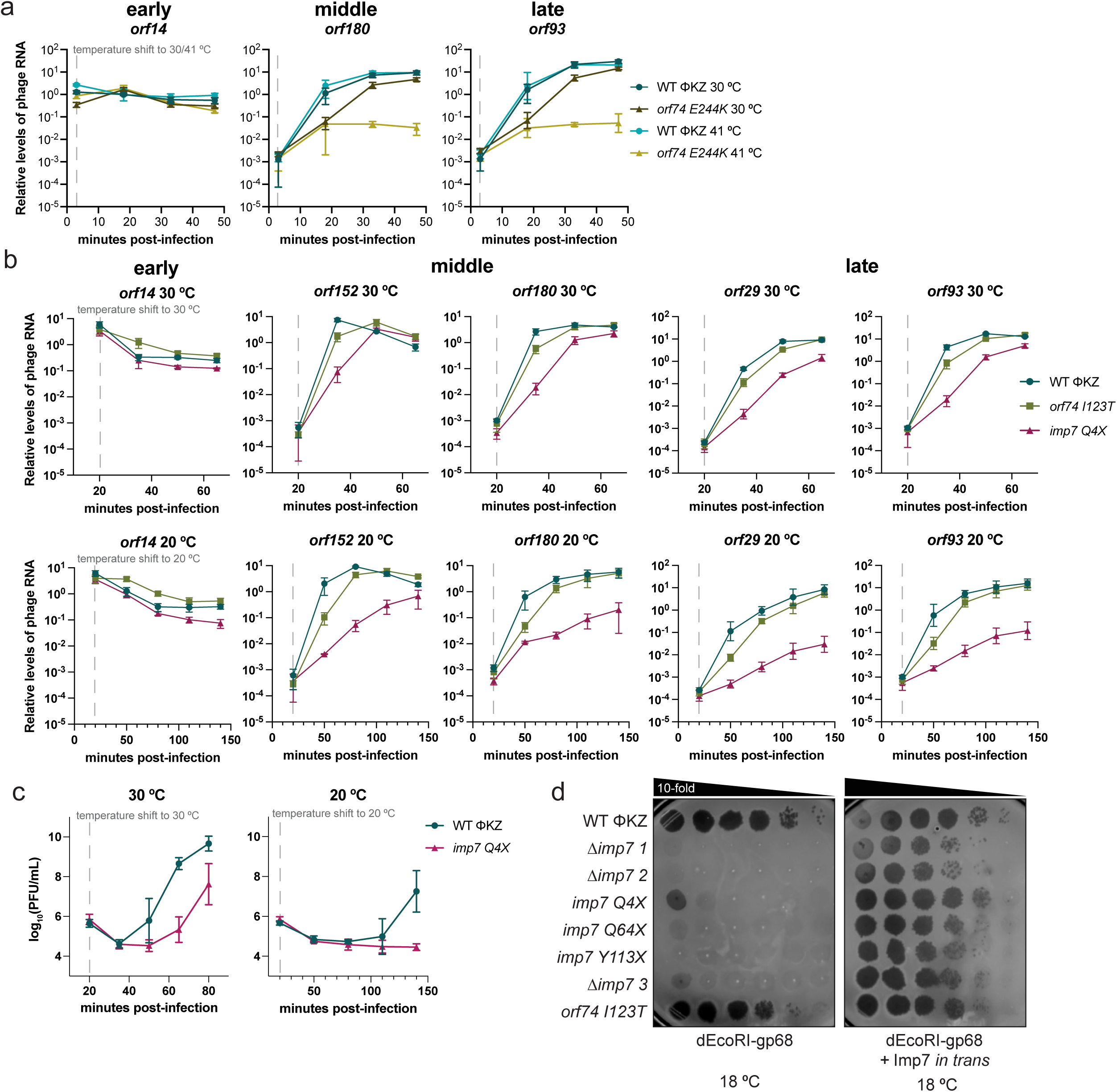
*imp7* is conditionally essential at environmental temperatures. a-b, RT-qPCR measuring phage transcript abundance over time relative to host transcript rpoD, using primers to amplify the indicated gene during infection at the indicated temperature. c, log_10_ PFU/mL output from 30 °C and 20 °C infections from b. d, Plaque assays on the indicated PAO1 strain, spotting 10-fold serial dilutions of the indicated wildtype or mutant phage at the indicated temperature. Transcript measurements, PFU/mL measurements, and plaque assays were performed in three independent biological replicates.

### Transcriptional output is tied to nvRNAP import efficiency

Imp7 is required for maximal import of the essential nvRNAP complex, but *imp7* nonsense mutations and deletions indicate that it is not strictly essential for phage replication in LB at 30 °C. We next questioned how phages were viable with decreased nvRNAP import, considering it is essential for phage replication^11,18^. We first assessed whether the bacterial RNAP becomes required by the phage when lacking *imp7*, by measuring mutant phage replication in the presence of host RNAP inhibitor rifampicin^7^. Rifampicin treatment decreased viable CFU by >5 orders of magnitude, but robust phage production was observed for wildtype and mutant phages (Sup Fig 1h). Therefore, a bacterium that has died due to host RNAP inhibition can still be a robust phage factory for WT and *imp7* mutant phages.

We next queried whether *imp7* mutants have less transcriptional output due to decreased nvRNAP in the phage nucleus. To assess the contribution of Imp7-dependent nvRNAP import to transcriptional output, we first confirmed that middle and late transcripts indeed depend on nvRNAP activity. Of note, it is presumed that the nvRNAP alone executes both middle and late transcription, although *in vitro* nvRNAP activity has only been reconstituted for late promoters^19,20^. We measured transcript levels for heat-sensitive *ts1 orf74 E244K* mutant phage over a single round of infection in liquid culture at 30 °C and 41 °C, the latter temperature being the non-permissive condition for this mutant. We found that both middle (*orf180*) and late (*orf93*)^7^ transcript levels were decreased by approximately 2 orders of magnitude after 45 minutes relative to transcript levels at 30 °C (Fig 2a). These data provide direct *in vivo* evidence that middle transcription depends on nvRNAP activity. We then used the same infection assay to measure the levels of middle and late transcripts in the *imp7 Q4X* mutant over one round of infection at 30 °C in liquid culture. The *imp7 Q4X* mutant displayed an initial delay in transcript production, producing approximately 2 orders of magnitude less transcript than wildtype, but by 65 minutes, the amount of these transcripts recovered to wildtype levels (Fig 2b). Consistent with this, we observed a 15-minute delay in burst time for the *imp7 Q4X* mutant phage relative to WT ΦKZ (Fig 2c). These data suggest that at 30 °C, wildtype phages have excess nvRNAP import capacity and thus even a large decrease in nvRNAP localization to the phage nucleus that lowers mRNA production does not abolish phage replication at 30 °C.

*imp7* loss-of-function mutants may be nonviable under conditions where nvRNAP import is more limited. We plated *imp7* mutants at high (42 °C) and a range of lowered temperatures (18-22 °C), mimicking environmental conditions. *imp7* mutants failed to plaque on solid agar at 18-20 °C while wildtype and nvRNAP mutant phages were unaffected (Fig 2d, Sup Fig 3a). Complementation with *imp7* expressed *in trans* restored plaquing at 18 °C, confirming that *imp7* is required for phage replication at environmental temperatures (Fig 2d). Strikingly, nvRNAP-dependent transcript levels in the *imp7* mutant phage at 20 °C were ∼2 orders of magnitude less compared to wildtype phage by 80 minutes post-infection (mpi) (Fig 2b). This transcriptional defect at lowered temperature mirrors what we observed in heat-sensitive mutant *ts1* (*orf74 E244K*) at 41 °C (Fig 2a), further suggesting that the *imp7 Q4X* mutant’s middle and late transcription defects are due to insufficient imported nvRNAP. Early (*orf14*) transcript levels were unaffected in the *imp7* mutant phage, and gp55 import-deficient *orf74 I123T* phage replicated well with normal transcription at 30 °C and 20 °C (Fig 2b). Together, these data demonstrate that nvRNAP import defects decrease middle and late transcriptional output, and illustrate the essential role *imp7* plays in nvRNAP import during growth at environmental temperatures.

**Figure 3:**
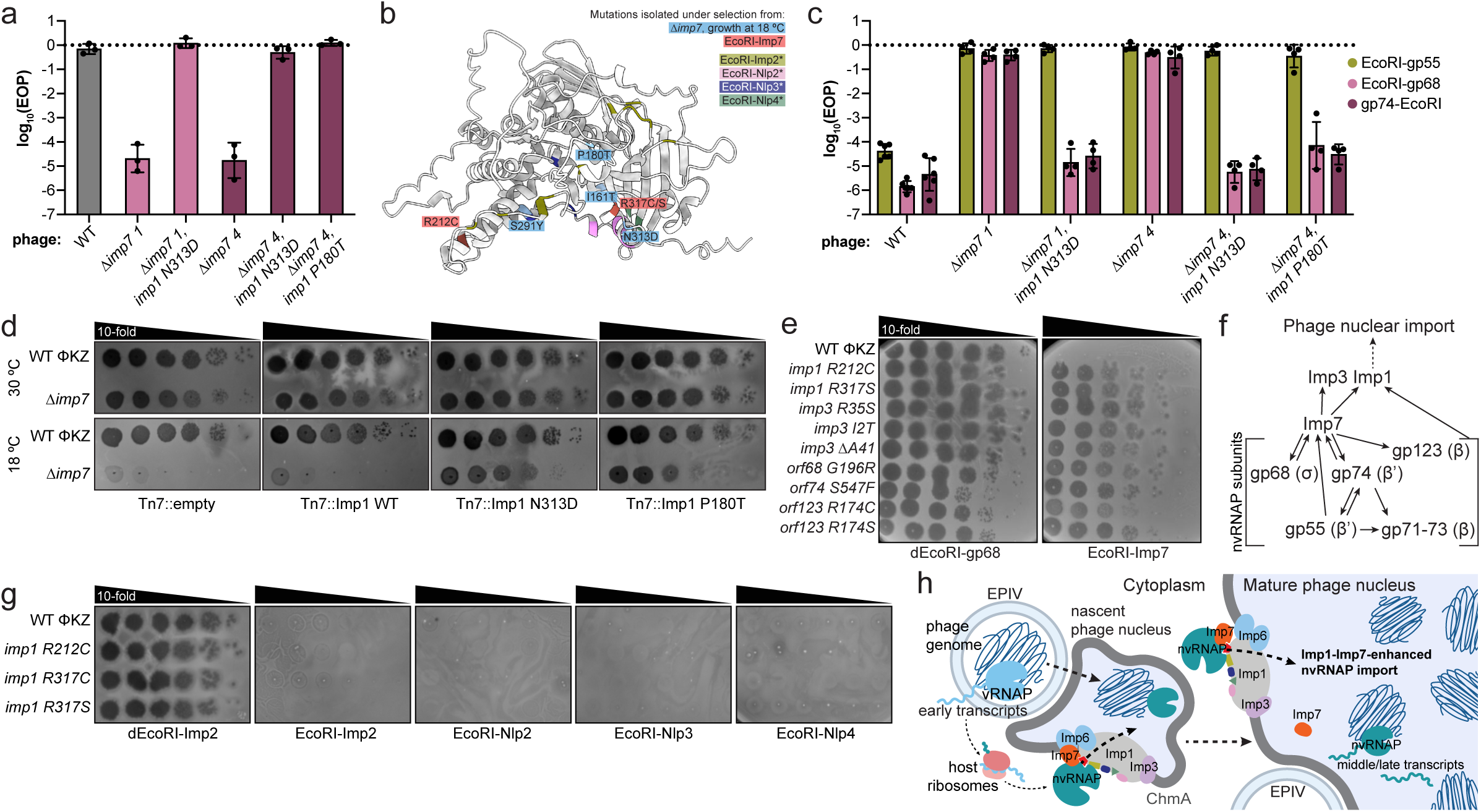
Imp1 is required for nvRNAP import. a, Log_10_ efficiency of plating (EOP) of wildtype or mutant phage on catalytically dead EcoRI-gp68 (dEcoRI-gp68) expressing strain at 18 °C relative to phage titer on the same strain at 30 °C. b, AlphaFold 2 structural model of Imp1 with amino acids mutated in various EcoRI-nvRNAP selections color coded. EcoRI-fusion* indicates previously identified Imp1 mutations isolated from selections in Kokontis *et al*.^13^, presented here for comparison. c, Log_10_ efficiency of plating (EOP) of wildtype or mutant phage on the indicated EcoRI-nvRNAP expressing strain at 30 °C, relative to phage titer on PAO1 strain expressing dEcoRI-gp68 at 30 °C. d, Plaque assays on PAO1 strain expressing the indicated Imp1 wildtype or suppressor mutant allele from the PAO1 *att*Tn7 chromosomal site, spotting 10-fold serial dilutions of wildtype or Δ*imp7::acrVIA1* replacement phage (Δ*imp7*) at the indicated temperature. e, g, Plaque assays on PAO1 strain expressing the indicated EcoRI-nvRNAP fusion, spotting 10-fold serial dilutions of the indicated wildtype or mutant phage at the indicated temperature. f, Graphical representation of genetic requirements for Imp7 and nvRNAP import. h, Model cartoon depicting Imp1-mediated and Imp7-enhanced import of nvRNAP into the phage nucleus. Plaque assays and efficiency of plating experiments were performed in at least three independent biological replicates with similar results.

### nvRNAP import is mediated by Imp1 and enhanced by Imp7

Phage nvRNAP import appears to operate in both an Imp7-dependent and an Imp7-independent manner. This Imp7-independent nvRNAP import is apparently active at 30 °C where *Δimp7* phages are viable and still execute middle/late transcription, albeit with a delay. To identify genes required for Imp7-independent nvRNAP import, we reasoned that we could “evolve” an improved version of Imp7-independent nvRNAP import at lowered environmental temperatures. Plating high titers of two phages (Δ*imp7 1* and Δ*imp7 4*, Fig 1b) at 18 °C where they are inviable, we isolated mutant suppressor phages that can grow at 18 °C by compensating for loss of *imp7*. Rare suppressor phages arose by acquiring spontaneous mutations, and whole genome sequencing revealed suppressor mutations in *imp1* (Table 1, Sup Table 1), at new *imp1* residues not previously characterized (Fig 3a, 3b, Sup Fig 3b). We reasoned that these suppressor alleles may restore or enhance nvRNAP import in the absence of Imp7, thereby restoring transcription and phage viability at 18 °C. To test this, we plated suppressor mutants on EcoRI-nvRNAP fusions and found that *Δimp7-imp1* suppressor mutant phages became resensitized to targeting by two of the three EcoRI-nvRNAP subunit fusions (Fig 3c), consistent with increased import of nvRNAP subunits in the absence of Imp7 to suppress lethality at 18 °C. Together, these data establish Imp1 as a component of the nvRNAP import pathway with maximal import enabled by Imp7. Finally, over-expression of *imp1 N313D* and *P180T* mutant alleles rescue Δ*imp7* phage plaquing at 18 °C when expressed *in trans* while wildtype *imp1* does not (Fig 3d). This inability of over-expressed wildtype Imp1 to enable import of sufficient quantities of nvRNAP in the absence of Imp7 suggests that Imp1 is not perfectly suited for nvRNAP import and needs either mutated residues or Imp7. Together, these data demonstrate that Imp1 is required for nvRNAP import, with maximal import enabled by Imp7, which is critical at environmental temperatures.

To understand whether Imp7 enhances Imp1’s nvRNAP import function or whether the two proteins operate independently from one another, we assessed whether Imp7 is imported into the phage nucleus (Sup Fig 3c). If Imp7 depends on Imp1 for its import, rather than an independent pathway, this suggests that Imp7 and Imp1 operate within the same pathway to import nvRNAP. We fused Imp7 to EcoRI to define its import requirements, and EcoRI-Imp7 fusions restricted wildtype phage by ∼10^7^-fold (Sup Fig 1a), and two classes of escape mutations emerged (Fig 3e, Table 1, Sup Table 1). First, mutations in the genes encoding the nvRNAP subunits *orf68*, *orf74*, and *orf123* decreased Imp7 import, suggesting that import of Imp7 and nvRNAP are co-dependent. The second class were mutations in known import factors *imp1* and its essential co-factor *imp3*. This latter result confirms that Imp7 operates within the established Imp1-dependent import pathway (Fig 3f). Expression of Imp1 *in trans* resensitized *imp1* mutant phages to EcoRI-Imp7 targeting, confirming these *imp1* mutations are causal for decreased EcoRI-Imp7 import (Sup Fig 3d). The Imp7-specific *imp1* mutations (*R212C and R317C/S*) are unique from any other Imp1 alleles identified by our previous selections^13^, suggesting a new interface for Imp7/nvRNAP import (Fig 3b, Sup Fig 3b). Consistent with this prediction, Imp7-selected *imp1* mutant phages still imported EcoRI fusions to *imp2*, gp155/Nlp2, gp104/Nlp3, and gp171/Nlp4, suggesting that this interface is indeed specific to Imp7 (Fig 3g). Together, we propose that Imp7 is an adaptation for ΦKZ, acting as an “enhancer” of Imp1’s intrinsic suboptimal nvRNAP import activity (Fig 3h).

### Phage nucleus assembly is uncoupled from DNA replication and middle transcription

To understand the impact of middle and late transcriptional defects on the progression of infection, we used live-cell fluorescence microscopy to track infection of WT, *imp7 Q4X*, and *Δimp7 3* mutant phages over time at 30 °C and 20 °C. Genome injection and migration of the injected genome to the center of the cell proceeded normally in *imp7* mutants at 20 °C, but the DAPI-stained genome remained small in ∼90% cells at 140 mpi (Fig 4a). This observation is similar to when phage DNA replication failure was induced by ciprofloxacin^14^. qPCR of a single round of infection at 20 °C confirmed that *imp7 Q4X* phages fail to replicate DNA without *imp7* (Fig 4b), where DNA replication is robust at 30 °C (Fig 4b), but with a slight delay that mirrors the delay in middle and late transcription (Fig 2b). Expression of Imp7 *in trans* restored DAPI-stained nucleus size for both *imp7* mutants, confirming replication failure is due to defects in nvRNAP-dependent transcription (Fig 4a).

**Figure 4:**
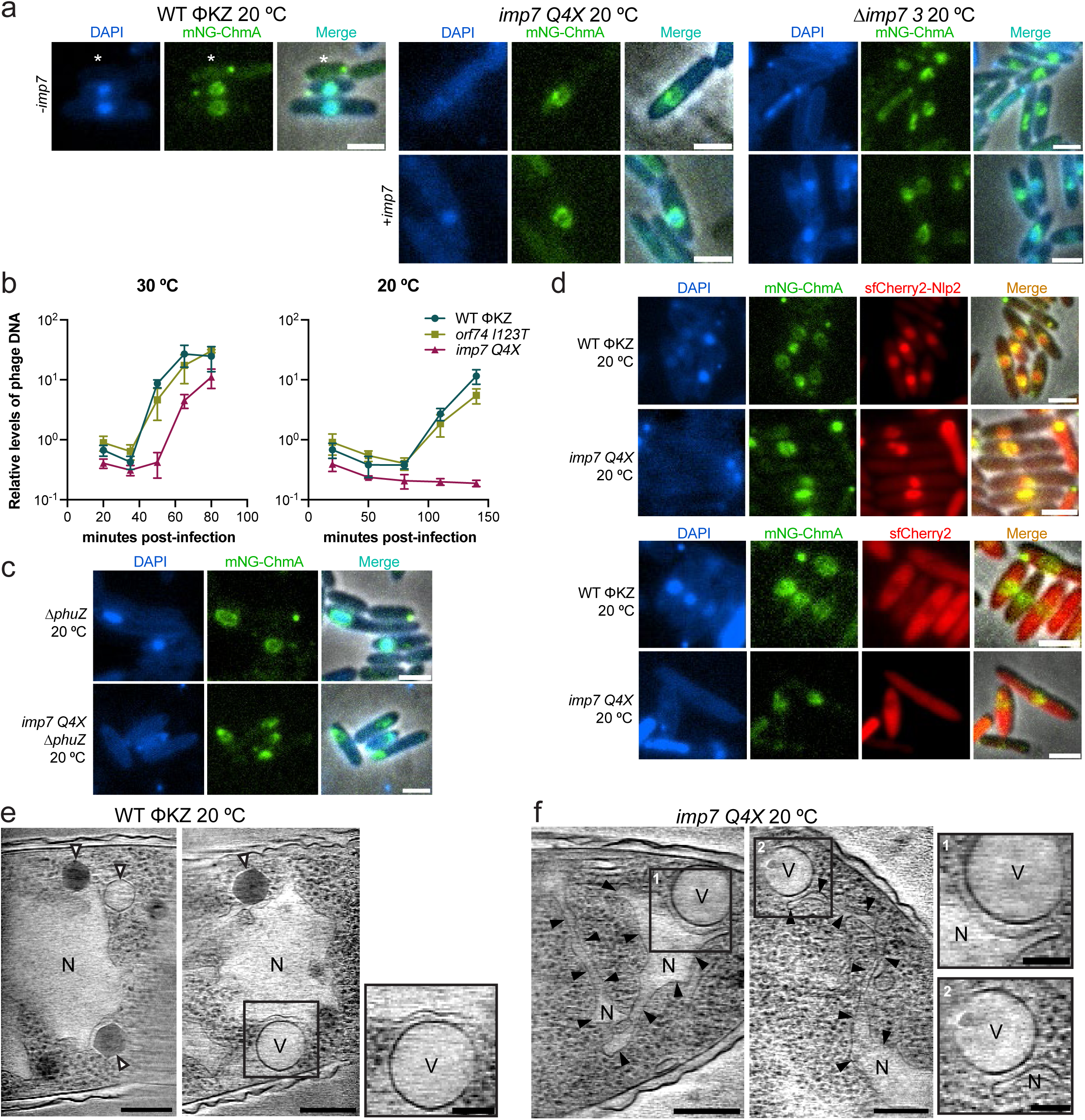
Phage nuclear shell assembly precedes DNA replication and middle/late transcription. a, Wildtype or indicated mutant phage infecting PAO1 strain expressing mNeonGreen-ChmA at 20 °C at 140 mpi, with or without Imp7 expression from the PAO1 *att*Tn7 chromosomal site (+/− *imp7*) (Without complementation, WT ΦKZ n=358 cells, *imp7 Q4X* n=693 cells, Δ*imp7 3* n=576 cells. With complementation, *imp7 Q4X* n=620 cells, Δ*imp7 3* n=527 cells). White asterisks indicate uninfected cells. b, Quantification of phage DNA replication via qPCR using primers for ChmA during infection at 30 °C or 20 °C, with the indicated wildtype or mutant phage. c, *ΔphuZ* or *imp7 Q4X, ΔphuZ* phage infecting PAO1 strain expressing mNeonGreen-ChmA at 20 °C at 140 mpi (*ΔphuZ* n=257 cells, *imp7 Q4X, ΔphuZ* n=559 cells). d, Wildtype or *imp7 Q4X* mutant phage infecting PAO1 strain expressing mNeonGreen-ChmA and sfCherry2-Nlp2 (WT ΦKZ: n=812 cells, *imp7 Q4X*: n=460 cells) or sfCherry2 (WT ΦKZ n=394 cells, *imp7 Q4X* n=100 cells with nuclear exclusion and n=134 cells with diffuse localization) at 20 °C at 140 mpi. e-f, Tomographic slices of wildtype and *imp7 Q4X* mutant phage infected PAO1 strain expressing mNeonGreen-ChmA at 20 °C at 140 mpi (two representative cells). White arrows, capsid. N, phage nucleus. V, vesicle. f, Black arrows indicate nuclear shell walls. Microscopy was performed at least two independent times in biological replicates with similar results. Single round infections for DNA measurements were performed in three independent biological replicates, with the last timepoint performed in two independent biological replicates. Cryo-electron tomography was performed once (WT ΦKZ) or twice (*imp7 Q4X*). Scale bars, 2 μm (a, c, d), 200nm (e, f), 100nm (inset e,f).

The *imp7 Q4X* mutant phage provides an ideal conditional essential mutant to determine which steps of the phage developmental cycle require DNA replication and middle transcription. When we visualized shell formation by infecting cells expressing mNeonGreen-ChmA (mNG-ChmA) fusions, we were surprised to see an mNG-ChmA structure assembling at the center of the cell despite the lack of replicating phage DNA (Fig 4a). This suggests that DNA replication is not required to drive or support ChmA oligomerization. Nascent nuclear compartments in *imp7 Q4X* infections often appeared smaller and less well-formed than wildtype ChmA compartments but were nonetheless ChmA assemblies. Expression of mNG-ChmA in uninfected cells does not result in similar structures of this apparent size or those that are centered in the cell (Fig 4a, asterisks), suggesting that early-expressed phage-encoded factors coordinate assembly of this large compartment *in vivo*, independently of middle transcription and DNA replication.

To assess whether ChmA structures are properly developing nucleus compartments or are spurious ChmA assemblies, we queried whether two key processes in phage development were occurring: PhuZ-dependent shell migration and phage nuclear protein import. Deletion of *phuZ* (ΦKZ *orf39*) in the *imp7 Q4X* background confirmed that midcell migration of the developing mNG-ChmA assembly is PhuZ-dependent (Fig 4c), suggesting that ChmA is in a suitable oligomeric state to interact with PhuZ. To determine whether mNG-ChmA is oligomerizing to form an enclosed compartment, even if devoid of replicating DNA, we assessed the localization of nuclear (sfCherry2-Nlp2) and cytoplasmic (sfCherry2) proteins at 20 °C. We observed concentrated sfCherry2-Nlp2 (Fig 4d), suggesting ChmA assemblies represent compartments with enclosed, interior space that are competent for protein import. sfCherry2 was generally excluded from these nuclear assemblies, while appearing diffuse in some cells (Fig 4d). Together, these observations demonstrate that closed, selectively permeable nuclear compartments can assemble without DNA replication and middle/late transcription.

Having defined a semi-functional ChmA compartment that imports/excludes select proteins and is trafficked to the mid-cell but lacks nvRNAP, middle transcripts, and replicating DNA, we next asked what the macrostructure of this assembly is. Using cryo-electron tomography (cryo-ET), we observed *in situ* phage nuclei during wildtype and *imp7 Q4X* infections at 20 °C at 140 mpi. Wildtype infections progressed to form large nuclei, often with packaged capsids visible or assembling virions nearby (Fig 4e, Sup Fig 4a-d, Supplementary Video 1). *imp7 Q4X* nuclear compartments were smaller and thinner, however, sometimes with clear interior space while other times appearing like a sheet folded on itself (Fig 4f, Sup Fig 4e-h, Supplementary Video 2). Notably, in both wildtype and *imp7 Q4X* infections, this compartment shows a clear difference in density on the interior from the cytoplasm and excludes host ribosomes, consistent with these ChmA assemblies being closed compartments. EPI vesicles were also clearly visible, but we did not observe capsid or virion assembly in arrested *imp7 Q4X* infections as expected, because late genes encoding structural proteins (*i.e.* portal gene *orf29*) are not transcribed in the absence of nvRNAP. Intriguingly, we did observe consistent and close contact between the EPI vesicle and the phage nucleus in both wildtype and *imp7 Q4X* arrested infections (Fig 4e-f inset, Sup Fig 4a-b, d-h). The factors and mechanisms facilitating this connection remain to be resolved. In sum, these data suggest that nvRNAP import/activity and subsequent DNA replication are needed to fully grow the ChmA phage nucleus but that it assembles and establishes cytoplasmic segregation prior to these activities. Phage infection, when arrested at this time point, could provide a valuable tool in downstream studies focused on EPI vesicle-nucleus interactions.

## Discussion

ΦKZ-like jumbo phages exhibit remarkable segregated subcellular organization during infection, where two selective compartments segregate phage transcription/DNA replication spatially and temporally. The transcriptional program is a clear example of this, where production of early transcripts is physically separated from the site of middle/late transcription, as the vRNAP is injected and remains inside the EPI vesicle at the start of infection. nvRNAP is an early product and is localized to a separate nucleus-like compartment to execute nvRNAP-dependent transcription. In this way, coordinated early-middle-late transcriptional programs with genome handoff from the EPI vesicle to the nucleus, and proper import of nvRNAP into the nucleus for function, are key developmental steps for replication.

Here we describe a tripartite import pathway that regulates nvRNAP import into the phage nucleus. Imp1 is required for nvRNAP import, and Imp1’s function is enhanced by Imp7, a novel import factor and previously uncharacterized gene. By sequence, Imp7 is not well-conserved beyond close ΦKZ relatives, suggesting that Imp7 may be an adaptation for ΦKZ and its conserved Imp1 protein to ensure import of a critical enzyme complex at a broad range of temperatures. In addition, we find that nvRNAP subunits are co-dependent for import, consistent with recognition and/or import of an assembled or partially assembled complex by Imp1/Imp7. How such a large (>300 kDa) complex is trafficked through the nuclear wall remains unknown.

We also find that defects in middle/late transcription caused an intermediate point of failure, after successful adsorption, injection, and early transcription. Some arrest mechanisms described previously, like early mRNA degradation^12^, translational arrest^3^, and EPI vesicle-sensing host immune systems^22^, act early and halt infection at the site of injection, before the nascent proteinaceous nucleus can be expressed or assembled. Middle transcriptional arrest provides a distinct window into the next critical transition period when the nascent nuclear shell is established, similar to ASO-mediated knockdown of essential middle genes like *orf155*/*nlp2*^11^. While we saw no phage DNA replication in the absence of middle transcripts as expected, we unexpectedly found that nuclear shell oligomerization proceeds without DNA replication or middle transcription. Although shell monomers have intrinsic affinity to oligomerize *in vitro*^10^, large assemblies (at the µm scale) are not observed in uninfected cells *in vivo*. This suggests that early phage gene products, potentially combined with host factors, are sufficient to initiate assembly of the nuclear shell into a functional selectively permeable compartment. Discovery of the factors required for selective macromolecule movement between compartments in this unique phage family will enable future biophysical investigation of this remarkably robust trafficking event.

## Acknowledgements

J.B.-D. is supported by the National Institutes of Health (nos. R01 AI171041 and R01 AI167412). C.K. received support from the National Institutes of Health (no. 2T32AI060537-21A1) and the UCSF Discovery Fellowship. D.M. received support from the National Institutes of Health Ruth L. Kirschstein National Research Service Award 1F32GM149125-01 and is currently supported by 1K99GM160780-01A1 (NIH-NIGMS). D.C. received support from the UCSF Discovery Fellowship. Cryo-EM equipment at UCSF is partially supported by National Institutes of Health grants S10OD020054, S10OD021741 and S10OD026881 and Howard Hughes Medical Institute. D.A.A. is supported by the Chan Zuckerberg Initiative. We thank Bondy-Denomy laboratory members for helpful insight and discussion. We additionally thank C. Gross for vital input. The nvRNAP 4 subunit expression vector was graciously shared by the Maria Yakunina lab. We thank Jingwen Guan for construction of the mNeonGreen-gp54 plasmid, and Li Yuping for generating the initial pool of HA-mutagenized ΦKZ.

## Supplemental information

Document S1: Figures S1-S4, Table S1

## Author contributions

C.K. – Conceptualization, Investigation, Visualization, Writing – original draft, Writing - review and editing

D.M. – Investigation, Writing – review and editing

D.C. – Investigation, Writing – review and editing

I.Z. – Investigation, Writing – review and editing

W.X. – Investigation, Writing – review and editing

D.B. – Investigation

D.A. – Funding acquisition, Writing – review and editing

J.B.D. - Conceptualization, Funding acquisition, Investigation, Writing - original draft, Writing - review and editing

---

**Sup Fig 1:**
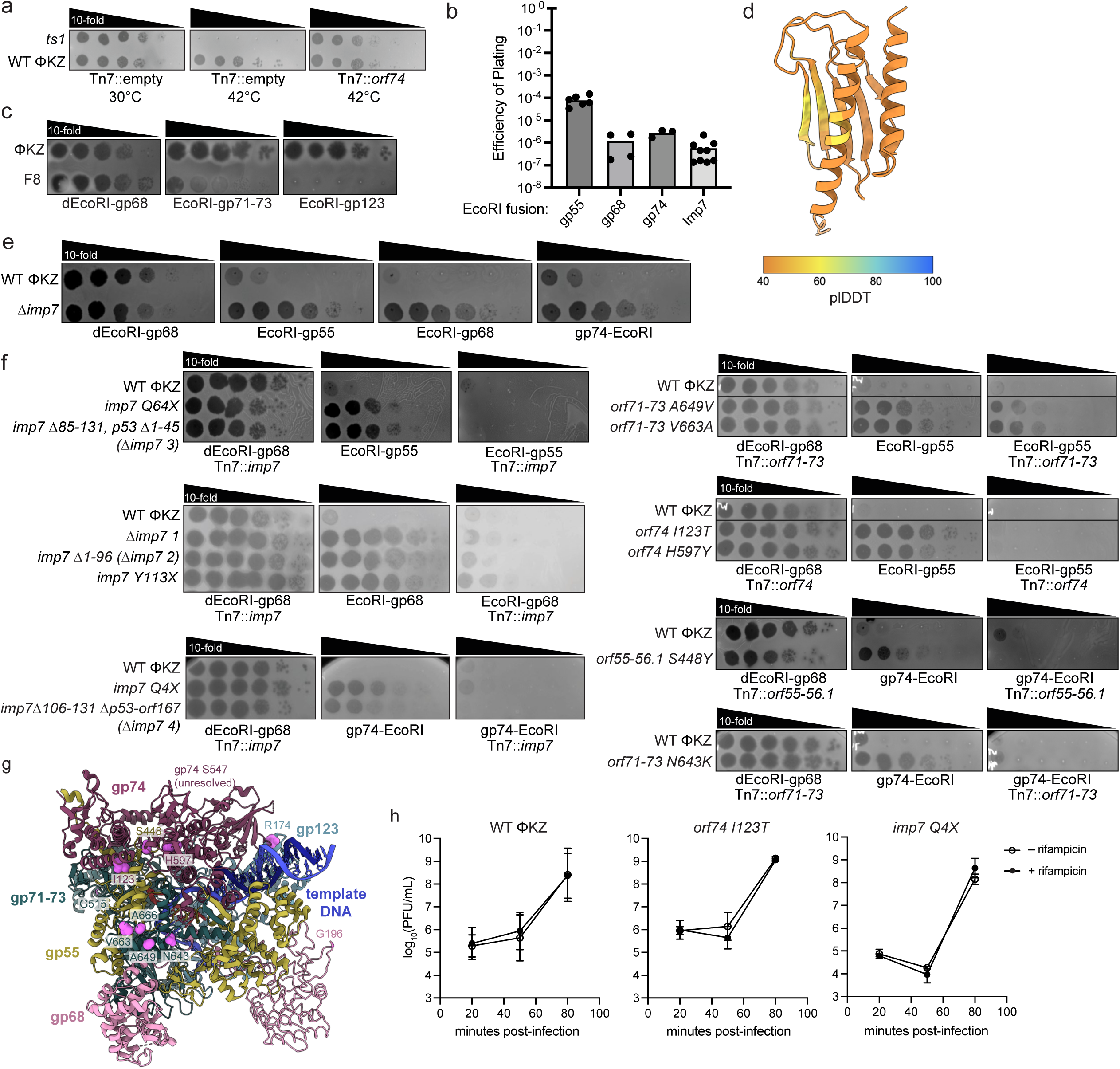
Mutations in *imp7* and nvRNAP subunits are causal to reduced nvRNAP import. a, Plaque assays of strain PAO1 with or without expression of *orf74* from the chromosomal *att*Tn7 site, spotting a 10-fold serial dilution of WT ΦKZ or *ts1 orf74* mutant phage, grown at 30 °C or 42 °C. b, Efficiency of plating of WT ΦKZ infecting PAO1 strain expressing the indicated EcoRI-nvRNAP fusion relative to PAO1 strain expressing catalytically dead EcoRI-gp68 (dEcoRI-gp68). c, Plaque assays of WT ΦKZ and F8 (EcoRI-sensitive negative control) phages spotted in 10-fold serial dilutions on lawns of PAO1 expressing the indicated EcoRI-nvRNAP fusion. d, AlphaFold 3 model of Imp7 colored by plDDT. e, Plaque assays of WT ΦKZ or *imp7::acrVIA1* replacement phage (Δ*imp7*) spotted in 10-fold serial dilution on a lawn of bacteria expressing the indicated EcoRI-nvRNAP fusion. f, Plaque assays of WT ΦKZ or indicated mutant phages spotted in 10-fold serial dilution on a lawn of bacteria expressing the indicated EcoRI-nvRNAP fusion, with or without complementation of the indicated gene *in trans* expressed from the chromosomal *att*Tn7 site in PAO1. g, nvRNAP structure bound to template DNA (PDB:8QUE^23^) with residues mutated under EcoRI selections labeled, colored in magenta and side chains modeled as spheres. h, Log_10_ PFU/mL titer output from single round infection experiments with or without rifampicin treatment. All plaque assays and single round of infection experiments were performed in independent biological triplicates with similar results.

**Sup Fig 2:**
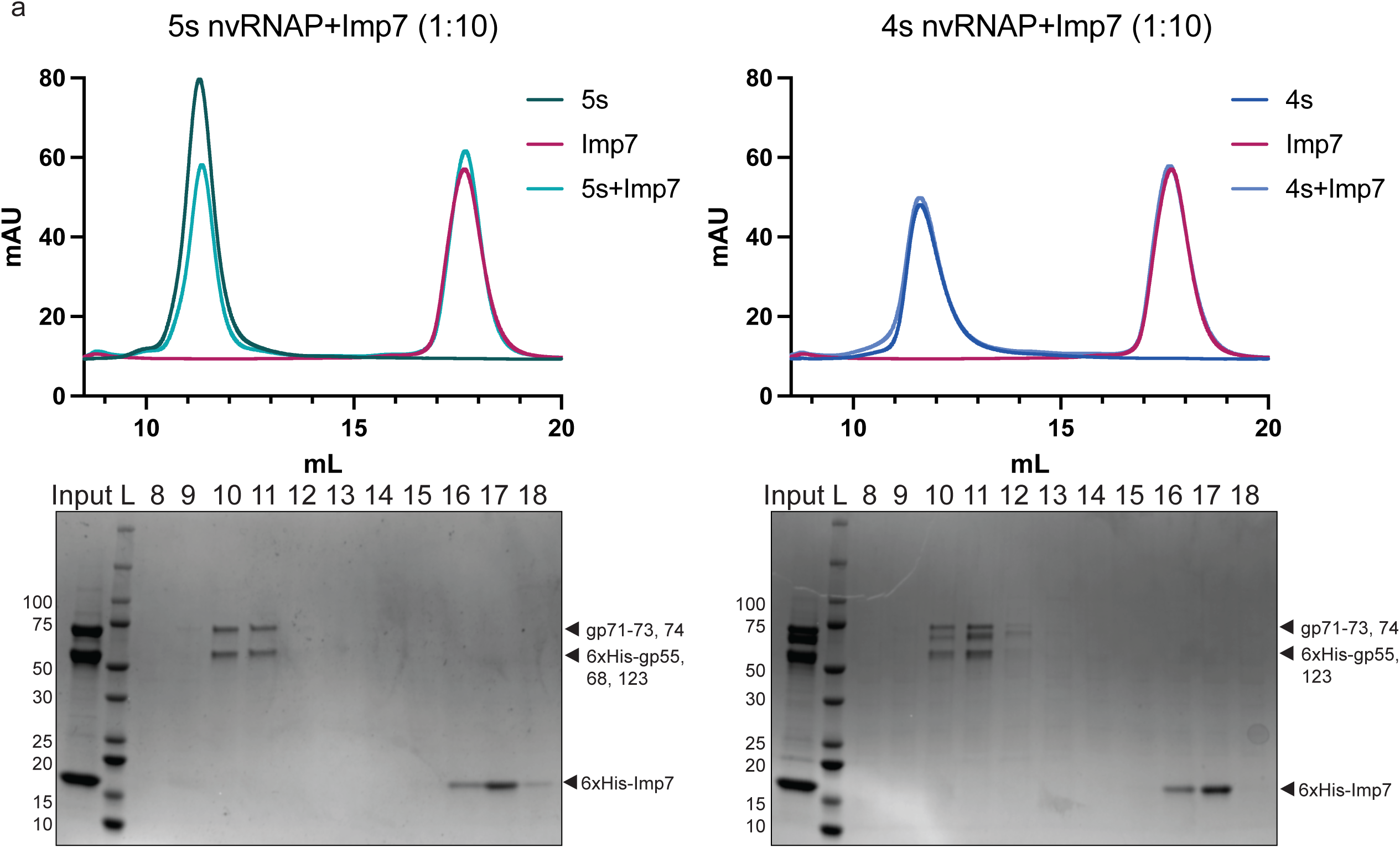
Imp7 does not bind nvRNAP *in vitro*. a, Size-exclusion chromatography trace of indicated protein mixtures, with corresponding fractions run on Coomassie-stained SDS PAGE gels. Proteins of interest are indicated with black arrows. Binding assays were performed two independent times with similar results.

**Sup Fig 3:**
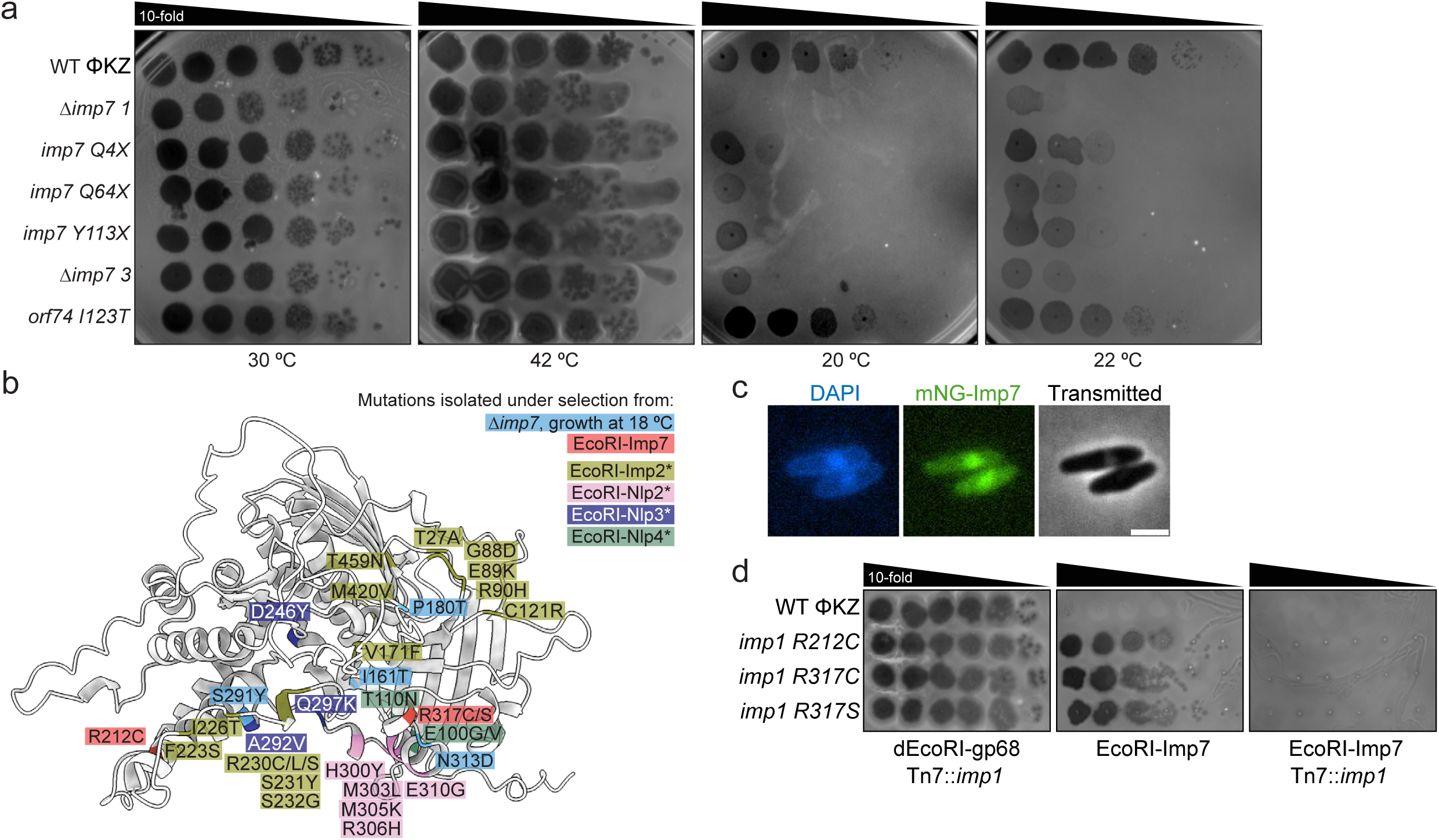
Imp7 is conditionally essential at a range of lower environmental temperatures. a, Plaque assays on the indicated PAO1 strain, spotting 10-fold serial dilutions of the indicated wildtype or mutant phage at the indicated temperature. b, AlphaFold 2 structural model of Imp1 with amino acids mutated under selection from various EcoRI fusions color coded. EcoRI-fusion* indicates previously identified Imp1 mutations isolated from selections in Kokontis *et al*.^13^, presented here for comparison. c, Live-cell fluorescence microscopy of WT ΦKZ infecting PAO1 strain expressing mNeonGreen (mNG)-Imp7 (n=322 cells) Scale bar, 2 µm. d, Plaque assays of WT ΦKZ or indicated mutant phages spotted in 10-fold serial dilution on a lawn of bacteria expressing the indicated EcoRI fusion, with or without *imp1* expressed *in trans* from the chromosomal *att*Tn7 site in PAO1. Plaque assays were performed at least two (a) or three (d) independent times with similar results. Microscopy was performed in three independent biological replicates with similar results.

**Sup Fig 4:**
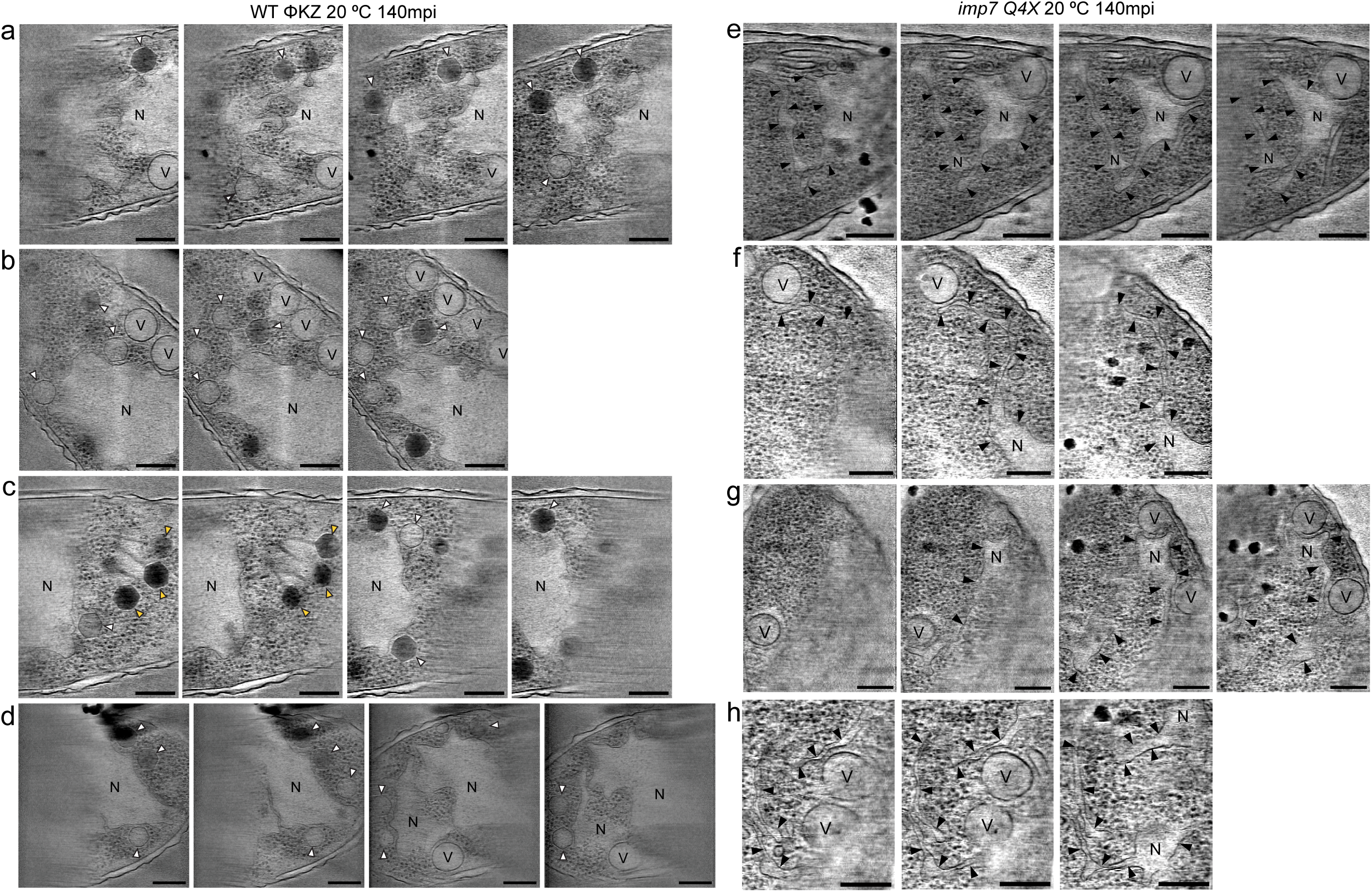
cryo-ET reveals arrested phage nuclei in the absence of nvRNAP. a-b, Tomographic slices of WT ΦKZ (a) or *imp7 Q4X* (b) infected PAO1 cells at 20 °C at 140 mpi. Each row represents a series through multiple slices of the same cell. White arrows, capsid. N, phage nucleus. V, vesicle. c, Yellow arrows indicate packaged and assembling virions. e-h, Black arrows indicate nuclear shell walls. Scale bars, 200 nm.

## Methods

### Bacterial growth

*P. aeruginosa* strain PAO1 and *E. coli* strains XL1Blue and BL21 (DE3) were grown in Lysogeny Broth (LB) at 37 °C shaking at 175 rpm. Bacteria were grown with appropriate antibiotics (gentamicin 50 µg/mL for all pHERD30T vectors, carbenicillin 200 µg/mL for pDuet vectors, kanamycin 50 µg/mL for nvRNAP-expressing vectors) and grown at 37 °C for 15-16 hr overnight shaking at 170 rpm. Bacteria were plated on solid LB agar including any necessary antibiotics for plasmid maintenance, and 10 mM MgSO_4_ was added when plating for phage infection. Protein expression from BL21 was induced with 1 mM IPTG. Expression of genes from PAO1 chromosomal *att*Tn7 site was induced with 1 mM IPTG in the LB solid agar. Basal expression of EcoRI-sfCherry2-nvRNAP or -Imp7 fusions from the leaky pHERD30T arabinose-inducible promoter was sufficient to restrict phage on plates, so L-arabinose was not included.

### Phage growth

Phages were grown by mixing 10 µL phage with 150 µL PAO1 overnight culture and 3.5 mL 0.35% top agar supplemented with 10 mM MgSO_4_, poured onto an LB agar plate containing 10 mM MgSO_4_, 50 µg/mL gentamicin, and any necessary inducers, and grown for 15-16hr overnight at 30 °C. The next day, phages were isolated by picking single plaques into 200 µL SM phage buffer. Escaper plaques were isolated in this way on restrictive strains, and purified three times by repeating this method. High titer lysates were generated by infecting PAO1 at the restrictive condition (either restrictive strain used in selection or restrictive temperature) overnight in LB 10mM MgSO_4_, with appropriate antibiotics and inducers at 37 °C (or the restrictive temperature). The supernatant was collected and treated with 5% volume chloroform, shaken gently for 10 min, and spun down for 5 min at max speed to remove cell debris. This was repeated 1-2 more times, and the final phage lysate was stored with 5% volume chloroform.

### Hot temperature sensitive phage isolation

Mutagenized phages were prepared via hydroxylamine (HA) chemical mutagenesis according to Villafane *et al*^24^. Briefly, 200 µL of ∼10^10–11^ PFU/mL WT ΦKZ was treated with HA by adding phage to a reaction buffer composed of 400 µL phosphate buffer (0.5 M KH_2_PO_4_, 5 mM EDTA, pH 6 with KOH), 400 µL ddH_2_O, 800 µL 1M HA stock solution (pH 6 with NaOH) for 48 hr at 37 °C to induce random point mutations. HA-treated phages were dialyzed at 4 °C in SM phage buffer with three changes of buffer, using 100 kDa MWCO Amicon spin filters (Millipore). Resulting mutagenized phage were plated on full plate infections with PAO1 as described above in Phage growth, and incubated overnight at 30 °C. The next day, two plates of 150 µL PAO1 overnight were poured and dried for phage replica plating at 30 °C and 41 °C. A water bath and thermometer were maintained in the 41 °C incubator to monitor internal temperature. Phages were replica plated onto two lawns of PAO1 by picking a single plaque with a toothpick and poking the surface of both plates in succession in the same location on either plate. Replica plates were incubated overnight at 30 °C or 41 °C, and resulting plaques were inspected for growth defects (i.e. plaque size) on the 41 °C plate compared to 30 °C, as well as compared to neighboring plaques to control for edge effects. Candidate temperature sensitive mutants that grew at 30 °C but failed to grow at 41 °C were picked from the 30 °C plate and stored in 200 µL SM phage buffer. Heat sensitive phage *ts1* was plaque purified and grown to high titer at 30 °C in liquid culture and whole genome sequenced to identify mutations.

### Construction of fusion proteins

The shuttle vector pHERD30T^25^ was used for cloning and expression of EcoRI-sfCherry2-nvRNAP subunit/Imp and mNeonGreen fusions in *P. aeruginosa* strain PAO1. For EcoRI-sfCherry2-gp fusions (C-terminal fusions to gp68, gp55-56.1, gp71-73, gp123, Imp7, referred to as EcoRI-gpX fusion for simplicity), the vector backbone was digested with SacI and SpeI restriction sites or PCR amplified and purified. For cloning ΦKZ genes, inserts were amplified by PCR from diluted phage lysates, and Hi-Fi Gibson Assembly (NEB) was used to join the fragments, according to the manufacturer’s instructions. The resulting assembly was transformed into XL1-Blue *E. coli* competent cells via heat shock at 42 °C for 30 seconds, with rescue growth in LB media at 37 °C for 30 min. EcoRI-sfCherry2 fusion to gp74 was generated by Genscript’s custom cloning service, and gp74 was cloned at the N-terminus of the fusion (i.e., gp74-EcoRI-sfCherry2) as the C-terminal fusion was inactive. The functional fusion is referred to as gp74-EcoRI for simplicity. All plasmid constructs Sanger sequenced (Quintara) over the insert using primer sets where at least one primer annealed to the vector backbone to check for accurate insertion, or were whole plasmid sequenced (Plasmidsaurus or Quintara). Verified plasmids were electroporated into competent PAO1 cells and selected on 50 ug/mL gentamicin.

For expression in BL21 *E. coli*, a plasmid containing the 4-subunit nvRNAP was obtained from the Yakunina Lab^19^. A plasmid containing the 5-subunit nvRNAP was constructed by adding gp68 to the 4-subunit plasmid by inserting a gp68 fragment with the restriction-digested plasmid via HiFi Gibson Assembly (NEB), following the manufacturer protocol. A T7 promoter and RBS were added upstream of gp68 using the NEB Q5 Site-Directed Mutagenesis kit, following the manufacturer protocol. 6xHis-Imp7 was cloned into pDuet via HiFi Gibson Assembly (NEB). Assembled vectors were transformed into XL1Blue competent cells via heat shock. Constructs were confirmed via Sanger sequencing and whole plasmid sequencing.

### Bacterial strain engineering

For integration of genes into the PAO1 *att*Tn7 site, genes of interest were cloned from the phage with overhangs and assembled into a digested miniTn7 vector at the SacI-PstI sites using Hifi Gibson Assembly (NEB) according to the manufacturer’s instructions. Plasmid sequences were confirmed by colony PCR of the multiple cloning site using primers to amplify the gene of interest from the plasmid backbone and sanger sequencing or whole plasmid sequencing. For integration into the PAO1 chromosomal *att*Tn7 site, competent PAO1 cells were transformed with assembled miniTn7 plasmid and helper plasmid pTNS3, following the protocol from Choi and Schweizer^26,27^. Successfully integrated strains were confirmed by PCR for presence of the integrated gentamicin cassette, as well as PCR for presence of the gene of interest in the genome using one internal primer and one genomic primer, as previously described^13,26^. To remove gentamicin cassettes from strains, pFlp2 was transformed to flip out the gentamicin cassette, yielding carbenicillin resistant and gentamicin sensitive colonies. pFlp2 plasmid was cured by plating on 5% sucrose plates^26^.

### Phage spot titration plaque assays

Plaque assays were performed by pouring a mixture of 150 µL PAO1 overnight culture and 3.5 mL 0.35% top agar onto an LB agar plate containing 10 mM MgSO_4_, 50 µg/mL gentamicin, and any necessary inducers (1 mM IPTG to induce expression from chromosomal *att*Tn7). Once solidified, 10-fold dilutions of phage were applied to the surface in 3 µL spots and allowed to dry. Plates were grown at 30 °C or 41-42 °C for 15-16 hr, or 20-22 °C for 24 hr. All plating at 18 °C was performed for 42-44 hr, with the exception of *Δimp7* mutant plating on PAO1 strains expressing *imp1* suppressor alleles from the *att*Tn7 site (Fig 3d), which were grown for 72 hr to see full development of complemented *imp7::acrVIA1 (“Δimp7*” phage) plaques. For plating performed at restrictive temperatures (18-20 °C, 41-42 °C), internal incubator temperature was confirmed by placing a thermometer in a water bath inside the incubator.

### Efficiency of plating

Efficiency of plating (EOP) was calculated by counting plaque forming units (PFU) from full plate infections or phage spot titration plaque assays, depending on the number of phage mutants and strains being surveyed. For full plate infections, 150 µL of PAO1 overnight culture was mixed with 5-10 µL phage at the appropriate dilution and then mixed with 3.5 mL of 0.35% top agar supplemented with 10 mM MgSO_4_ and poured on LB agar plates. Plates were incubated at 30 °C overnight, and plaques were counted the next day. Efficiency of plating (EOP) was calculated as the number of PFU/mL on the EcoRI targeting fusion relative to the number of PFU/mL on a nontargeting strain. EOP graphs were generated in Prism (v11.0.0).

### Whole genome sequencing

Phage genomes were sequenced via next generation sequencing (NGS) or nanopore sequencing (Angstrom Innovation). Phage genomic DNA was extracted from 200 µL of phage lysates by adding 200 µL of “2x phage lysis buffer” (final 1x concentration: 10 mM Tris, 10 mM EDTA, 100 ug/mL Proteinase K, 100 ug/mL RNase A, 0.5% SDS) to 200 µL high titer phage lysate (>10^9^ PFU/mL). This was incubated at 37 °C for 30 min, then at 55° C for 30min. Genomic DNA was purified either by phenol chloroform extraction and ethanol precipitation, or by using the DNA Clean & Concentrator Kit or Genomic DNA Clean & Concentrator Kit (Zymo Research). For NGS, 50-100 ng genomic DNA input was used to prepare whole genome sequencing libraries using the Illumina DNA Prep Kit. A modified protocol was used, with 5x reduced quantities of tagmentation reagents for each prep, except the bead washing step where the recommended quantities of Tagment Wash Buffer were used. PCR indexing-amplification was performed on-bead with Phusion Master Mix (NEB), using primers (IDT) that matched the Illumina DNA Prep Kit sequences. PCR reactions were amplified for 9-12 cycles, resolved by gel electrophoresis, and products in the 400 bp size range were excised from the gel and purified using the Gel DNA Recovery Kit (Zymo Research). Samples and pooled libraries were quantified by Qubit. Libraries were sequenced on the Illumina MiSeq using 150-cycle v3 reagents (for single end sequencing: 150 cycles, Read 1; 8 cycles, Index 1; 8 cycles, Index 2. For paired end sequencing: 75 cycles, Read 1; 8 cycles, Index 1; 8 cycles, Index 2, 75 cycles, Read 2). Data were demultiplexed on-instrument and trimmed using cutadapt (v1.15) to remove Illumina adapters. Trimmed reads were mapped using Bowtie 2.0^28^ (–very-sensitive-local alignments). Alignments were visualized using IGV (v2.19.1). Mutations were called if they were present in >90% sequencing reads at loci with at least 10x coverage.

### orf*::acrVIA1* replacement phages

*acrVIA1* was cloned into the pHERD30T dual crRNA/Homology arm plasmid^18^, including 500 bp homology arms upstream and downstream of *imp7* or *phuZ* to facilitate homologous recombination. PAO1 transformed with the appropriate *orf::acrVIA1* replacement plasmid and expressing *Lse* Cas13a from the chromosomal *att*Tn7 site^18^ were grown to OD600 of 0.5 in LB media supplemented with 10mM MgSO_4_, 50 µg/mL gentamicin, 1 mM IPTG, and 0.3% L-arabinose at 37 °C. Once at OD600 0.5, cultures were diluted 1:100 and infected with WT ΦKZ at a multiplicity of ∼10^−3^, and infections continued overnight. The next day, cultures were pelleted to remove cell debris, and the supernatant was treated with 5% volume chloroform at least 2 times. 10-50 µL of resulting phage lysates were plated in full plate infections with 150-300 µL of PAO1 *att*Tn7::*cas13aLse* transformed strain transformed with pHERD30T dual targeting crRNA guides^18^ on LB agar plates supplemented with 10 mM MgSO_4_, 50 µg/mL gentamicin, 1 mM IPTG, and 0.3% L-arabinose to isolate single plaques. The next day, resulting candidate *Lse* Cas13a-resistant plaques were picked and resuspended in SM buffer, representing candidate *orf::acrVIA1* replacement phages. Candidate *orf::acrVIA1, Lse* Cas13a-resistant phages were plaque purified three times on the *Lse* Cas13a dual crRNA guide selection strain, and *orf::acrVIA1* replacement was validated by PCR using phage genomic primers outside the homology arms to confirm the larger product representing insertion of the larger *acrVIA1* gene. High titer lysates were generated as described in Phage Growth.

### Single round infection assay

For imp7 mutant infections, PAO1 strain expressing dEcoRI-sfCherry-gp68 was diluted 1:100 from overnight culture into LB media supplemented with 10 mM MgSO_4_ and 50 µg/mL gentamicin and grown shaking at 170 rpm at 37 °C until an OD600 of 0.4. To maximize RNA yield, 8 mL cells were concentrated to 1 mL for each infection condition, and phage were added at an MOI of ∼1. Phages adsorbed for 2 min before spinning down at 10k x g for 1 min to wash off excess phage with LB media supplemented with 10mM MgSO_4_, 50 µg/mL gentamicin. The wash was repeated a total of two times. Infections were then diluted by 20x to distribute into 2 mL per infection timepoint and placed in an air-controlled incubator to begin the time course experiment. 500 µL of infected cells were collected for RNA or DNA extraction (see Phage DNA extraction or Phage RNA extraction methods) by spinning down at 10k x g for 1 min at 4 °C. The supernatant was removed and pellets were flash frozen for subsequent processing. For burst measurements, 500 µL of infected cells were mixed with 10% volume chloroform, shaken for 20 min, spun down to pellet chloroform and cell debris, and supernatants were diluted for spot assays.

For hot temperature sensitive infections, PAO1 strain expressing empty pHERD30T vector was diluted 1:100 from overnight culture into LB media supplemented with 10 mM MgSO_4_ and 50 µg/mL gentamicin and grown shaking at 170 rpm at 37 °C until an OD600 of 0.4. This subculture was then split in two, with half shaking for an additional half hour at 30 °C, and the other half shaking for the same time at 41 °C. To ensure that infections in the high restrictive temperature maintained the high temperature throughout infection, phages were added to cultures at an MOI of ∼1, adsorbed on the bench for 2 min, and placed in the appropriate incubator without additional washing steps. 200 µL aliquots of infected cells were removed for downstream RNA extraction and were pelleted and flash frozen as described above.

### Rifampicin infection assay

PAO1 cells transformed with empty pHERD30T vector were subcultured at 1:100 dilution from an overnight culture to an OD600 of 0.4 in LB media supplemented with 10 mM MgSO_4_ and 50 µg/mL gentamicin and grown shaking at 170 rpm at 37 °C until an OD600 of 0.4. A stock solution of 50 mg/mL rifampicin was made in DMSO, and rifampicin was added to rifampicin-treated cultures at a final concentration of 400 µg/mL (and an equal volume of DMSO was added to the untreated cultures), following Ceyssens *et al.*^7^ Cultures were treated with rifampicin for 5 min, and then phage was added at an MOI of ∼1 and allowed to adsorb for 2 min. Infections were washed twice in LB with 10 mM MgSO_4_ to wash off unadsorbed phage by spinning at 10k x g for 1 min. After the final wash, pelleted infections were resuspended in LB with 10 mM MgSO_4_ +/− 400 µg/mL rifampicin (DMSO for mock treatment), then shaken in a 30 °C water bath at 170 rpm for 60 min. To measure PFU/mL, 100 µL aliquots of infected cultures were mixed with 20 µL of chloroform at the indicated time point and shaken for at least 20 min. Phage outputs were 10-fold serially diluted and spot titration plaque assays were performed to calculate output titer.

### Phage DNA measurement by qPCR

500 µL aliquots from infection timepoints were spun down at 10k x g for 1 min, the supernatant was removed, and the pellet was flash frozen in liquid nitrogen and stored at −80 °C. DNA was extracted by thawing and resuspending pellets first in 100 µL water, then mixing with 100 µL of “2x phage lysis buffer” (see whole genome sequencing method), supplemented with RNase H and DNase, and heated at 37 °C for 30 min, then 55 °C for 30 min. Genomic DNA was extracted using the Genomic DNA Clean & Concentrator kit (Zymo) and quantified by nanodrop. DNA was diluted to 1 ng/µL for input to qPCR. Primers for *orf54* (Fwd: 5’-AAAGAAGAAGCAATCCGCC-3’, Rev: 5’-CAATGAGTAGACCAGCCATACC-3’) were used to amplify phage DNA, and primers for host *rpoD* (Fwd: 5’-GGGCGAAGAAGGAAATGGT-3’, Rev: 5’-CTGGATCAGGTCGAGGAATTG-3’) were used to amplify host DNA as a control. qPCR master mix (NEB Luna Universal qPCR master mix) was prepared according to the manufacturer’s protocol.

### Phage RNA measurement by RT-qPCR

500 µL aliquots from infection timepoints were spun down at 10k x g for 1 min, the supernatant was removed, and the pellet was flash frozen in liquid nitrogen and stored at −80 °C. RNA was extracted by resuspending pellets in 500 µL RNA lysis buffer (40 mM NaOAc, 1% SDS, 2mM EDTA). Then 500 µL pre-warmed (65 °C) acid phenol chloroform was added, vortexed to mix, and incubated in a 65 °C water bath for 5 min with frequent vortexing. Mixtures were then transferred to pre-spun phase-lock heavy tubes (5PRIME) and spun at 12k x g for 10 min. The aqueous layer was extracted at least twice with chloroform, or until the interface was clean. RNA was extracted via ethanol precipitation of the final aqueous layer. Total RNA was quantified by nanodrop. 10 µg total RNA was DNase I treated using the Turbo DNase kit (Thermo Fisher) with DNase I inactivation. DNase-treated RNA was quantified and diluted to 1 ng/µL for RT-qPCR input. RT-qPCR master mix (NEB Luna Universal One-Step RT-qPCR kit) was prepared according to the manufacturer’s protocol. For early phage transcripts, *orf14* primers were used (Fwd: 5’-gggttatcagccgtcgtaaa-3’, Rev: 5’-gggagcaggattcgaagatag-3’). For middle phage transcripts, *orf152* (Fwd: 5’-gatagcagctagctcaggataag-3’, Rev: 5’-gcacatgcaatgacttacgatac-3’) and *orf180* (Fwd: 5’-tcaaacacttcaccaactacac-3’, Rev: 5’-cgaccaacattcaaagatgcc-3’) primers were used. For late phage transcripts, *orf29* (Fwd: 5’-gtccgctattacgtctcttcac-3’, Rev: 5’-caagcgatgcttgaccataga-3’) and *orf93* (Fwd: 5’-acaagctaagatggctgaagat-3’, Rev: 5’-cccacttattcacacctgactta-3’) primers were used. For normalization control, host *rpoD* primers were used (see above, qPCR primers).

### Live-cell fluorescence microscopy

For EcoRI-sfCherry2-nvRNAP fusion and mNeonGreen-Imp7 fusion imaging, 0.8% LB agar pads (25% LB, 2.5 mM MgSO_4_) were supplemented with 1 µg/mL DAPI for phage DNA staining. PAO1 strains expressing the fusion construct from pHERD30T were subcultured at a 1:100 dilution with appropriate antibiotics and inducers (10 mM MgSO_4_, 50 µg/mL gentamicin, 0.01% L-arabinose induction for EcoRI-sfCherry2 fusions, 0.1% L-arabinose for induction of mNeonGreen-Imp7). Cells were grown until an OD600 ∼0.5 and then infected with phage for 60-80 min at 30 °C before imaging. Microscopy was performed on an inverted epifluorescence microscope (Ti2-E, Nikon) using the Perfect Focus System (PFS) and a Photometrics Prime 95B 25 mm camera. Cells were imaged through channels of phase contrast (200 ms exposure, for cell imaging), blue (DAPI, 50 ms exposure), green (FITC, 200 ms exposure, for mNeonGreen fusions), red (Cherry, 200 ms exposure, for sfCherry2 fusions) at 100x objective magnification (numerical aperture: 1.450).

For 20 °C imaging of *imp7* mutant phages, cells were grown and infected on solid media, similar to that described previously.^10^ Briefly, 1.2% LB agar pads (25% LB, 2.5 mM MgSO_4_) were prepared by solidifying 0.5 mL of melted LB agar in a deep well slide, supplemented with 1 µg/mL DAPI and appropriate inducers (1 mM IPTG for expression of Imp7 *in trans*, 0.1 mM IPTG for expression of sfCherry2 and sfCherry2-Nlp, or 0.1% L-arabinose for expression of mNeonGreen-gp54). A coverslip was placed on top to seal the well during solidification. Once solid, 5 colonies (grown from a streak plate the night before) of the indicated PAO1 strain were picked and resuspended in 100 µL of 25% LB MgSO_4_, and 5 µL colony suspension was spread onto the solidified agar pad. Inoculated slides were placed in a damp chamber at 30 °C and grown for 3-3.5 hr. 5 µL of high titer phage lysate (∼1E9-10 PFU/mL) was then spread on the slide. To mimic single round liquid infection conditions for PFU/DNA/RNA measurements, infected slides were incubated on the bench for 20 min (representing the washing time before time point samples begin). Slides were then grown for 120 min in a damp chamber in a 20 °C incubator before imaging (total infection time = 140 min). Final figure images were prepared in Fiji (version 2.16.0/1.54p).^29^

### Protein expression and purification

Plasmids expressing 4-subunit, 5-subunit nvRNAP (gp55 6xHis-tagged), or 6xHis-Imp7 were transformed and expressed in BL21 *E. coli.* Large scale subcultures were grown in LB media with appropriate antibiotics at 37 °C to an approximate OD600 of 0.5-0.7, starting from 1:100 dilution of overnight culture. Protein expression was induced with 1 mM IPTG at 18 °C for 16-18 hr. After growth, cells were pelleted at 4k x g for 20 min at 4 °C and resuspended in 40 mM Tris pH 8, 500 mM NaCl, 5 mM imidazole, supplemented with 1mM DTT, 1 tablet of protease inhibitor (EDTA free, Roche) and 25 U/mL Pierce Universal Nuclease (Thermo Fisher). Cells were lysed by sonication (15 s on, 45 s off, for 20 rounds at 50% amplitude). Lysates were clarified by spinning down at 30k x g for 30 min at 4 °C. Clarified lysate was removed from pellets and incubated with Ni-NTA resin (Qiagen) for 1 hr at 4 °C, using between 500 µL to 1 mL of resin bed volume per liter culture. The flow through was drained and the resin was washed with 100 bed volumes of Ni wash buffer (40 mM Tris pH 8, 500 mM NaCl, 20 mM imidazole, 1 mM DTT). Bound proteins were eluted by adding 5-10 bed volumes of Ni elution (40 mM Tris pH 8, 500 mM NaCl, 400 mM imidazole, 1 mM DTT) for 5 min. The elution was repeated twice. Elutions were concentrated to 500 µL before loading onto size exclusion chromatography by spinning in Amicon centrifugal concentrators. Concentrated elutions were loaded onto a Superdex 200 Increase 10/300 GL (Cytiva) and eluted with 40 mM Tris pH 8, 200 mM NaCl, 1 mM DTT. Fractions were flash frozen in liquid nitrogen and stored at −80 °C.

### Size exclusion chromatography binding assay

Purified proteins were mixed in the indicated molar ratios and incubated on ice for 30 min in a total volume of 500 µL. Protein complexes were loaded onto a Superdex 200 Increase 10/300 GL and peak fractions were analyzed by SDS-PAGE.

### Cryo-electron tomography sample preparation

50 mL of PAO1 expressing p30T-mNeonGreen-ChmA were grown for each infection by subculturing at 1:100 dilution from an overnight culture at 37 °C at 170 rpm in LB supplemented with 10 mM MgSO_4_, 50 µg/mL gentamicin, and 0.1% arabinose until OD600 reached ∼0.5. Once cultures were cooled to 20 °C, wildtype or *imp7 Q4X* phage were added at high titer such that most cells were infected with an effective multiplicity of infection (MOI) ∼1. Effective MOI was assessed via fluorescence microscopy beforehand by confirming that an injected genome was visualized at the pole (by DAPI staining of injected genomes and JukA localization to the EPI vesicle^22^). Infection proceeded on the benchtop for 20 min to mimic single round infection conditions performed for RNA/DNA/PFU measurements and 20 °C microscopy (described above). Infected cultures were then moved to 20 °C to shake at 170 rpm for 120 min. Cultures were then spun at 8k x g for 3 min to pellet infected cells, washed in 500 µL 50% bovine serum albumin (BSA) (wildtype and *imp7 Q4X* replicate 1) or 20% BSA (*imp7 Q4X* replicate 2) in phosphate-buffered saline (PBS), spun again at 10k x g for 2 min to pellet, and resuspended in 50 µL 50% BSA (wildtype and *imp7 Q4X* replicate 1) or 25 µL 20% BSA (*imp7 Q4X* replicate 2) and placed on ice before spotting on grids. *imp7 Q4X* replicates appeared similar regardless of differences in sample preparation.

#### High pressure freezing

5-7 µL of samples were applied onto the copper side of Formvar 200 mesh copper grids (TED Pella Inc., # 01700-F) freshly pre-coated with carbon (∼20 nm thick, TED Pella, #93010, Pointed Carbon Rods, Double Neck Tip 6.2 mm). The sample coated grids were vitrified in liquid nitrogen with the Leica EM-ICE High Pressure Freezer. Sample containing grids were high pressure vitrified between pre-polished and 1-hexadecence (Tokyo Chemical Industry #H0610) coated 6 mm PELCO Freezer hats, Brass specimen holder planchettes (TED Pella Inc., #39203). Grids containing vitrified cell samples were mounted into notched FIB Autogrid rings (nanosoft, #11021003) compatible with cryo-focused ion-beam (cryo-FIB) milling.

#### Milling

Samples were loaded into an Aquilos 2 dual beam microscope (FIB: Focused ion beam, SEM: Scanning electron microscope, Thermo Fisher Scientific) using a 35° Autogrid shuttle (copper bar side facing up, notch at 12 o’clock) to generate lamellae approximately 100-120 nm thick using the waffle method of milling^30,31^. Briefly, in the Aquilos chamber, the grid surface (copper side) was precoated sequentially with a platinum sputter coat (30 mA, 15 s, sputter vacuum), GIS (2 minutes, high vacuum), and a second platinum sputter coat (30 mA, 15 s, sputter vacuum). Thereafter, the grid was mapped using the Maps 3.28 software (Thermo Fisher Scientific) to identify potential lamellae sites. Thereafter, the grids were rotated 180° relative to mapping and tilted to a milling angle of 90° and trenches were milled with a 16 nA current (at regions approximately in the middle of a 200-mesh grid bar rectangle. Trench specifications: Dimension – Large rectangle: X = 22 µm, Y = 37 µm, Z = 1 µm; Small rectangle: X = 20 µm, Y = 17 µm, Z = 1 µm; Rectangle separation distance – 25 µm; overlap: X = 50 %, Y = 50 %; Scan Type – Serpentine; Scan Direction – Large Rectangle: Top to Bottom; Small Rectangle: Bottom to Top. After this, the grids were rotated back to mapping orientation, and a second grid map was recorded and lamellae sites were defined in the Maps software. The Maps project with defined lamella sites was loaded in AutoTEM Cryo 2.4. Thereafter the ‘Eucentric Tilt’ (Maximal Tilt Step = 10°; Preparation HFW = 250.0 µm, Resolution (guided) = 1536 × 1024) and ‘Milling Angle’ (Target milling angle = 20°; Clearance Angle = 2.0°; HFW = 160.0 µm) steps were performed. At each lamellae position, the bottom section of the lamellae site (large rectangle in trench) was further cleared by milling with an arbitrary rectangular milling pattern (Milling angle, Milling current: 45°, 7 nA; 35°, 7 nA; 20°: 3 nA). Thereafter a single notched stress relief cut was milled with the notch pointing outwards from the target lamellae site (Milling angle = 20°, Milling current = 0.3 nA). Thereafter the Image Acquisition (Ion HFW Oversize = 120%; Resolution: 1536 × 1024 at 4 µs, Enable ACB, Enable Auto Focus) and Lamellae placement (Ion HFW Oversize = 160%, Minimal Ion HFW = 50.0 µm) steps were executed with a target lamella dimension (12 µm × 10 µm × 100-120 nm). Lamellae were placed in alignment with the notch of the stress relief cut. Thereafter the ‘Milling’ and ‘Thinning’ steps were executed step-wise for each lamellae site. Milling details: Rough Milling (Pattern Offset = 1.0µm, Depth Correction, 100%, Front Width Overlap = 1.5 µm, Rear Width Overlap = 1.0 µm, Milling current = 1 nA, Pattern Type = CleaningCrossSection, DCM Rescan Interval = 120 s), Medium Milling (Pattern Offset = 800.0 nm, Overtilt = 0.5°, Depth Correction, 150%, Front Width Overlap = 650.0 nm, Rear Width Overlap = 500.0 nm, Milling current = 1 nA, Pattern Type = CleaningCrossSection, Pattern Overlap = 400.0 %, DCM Rescan Interval = 90 s), Fine Milling (Pattern Offset = 600.0 nm, Overtilt = 0.3°, Depth Correction, 150 %, Front Width Overlap = 350.0 nm, Rear Width Overlap = 100.0 nm, Milling current = 0.5 nA, Pattern Type = CleaningCrossSection, Pattern Overlap = 200.0%, DCM Rescan Interval = 60 s), Finer Milling (Pattern Offset = 400.0 nm, Overtilt = 0.1°, Depth Correction, 150 %, Front Width Overlap = 50.0 nm, Rear Width Overlap = 50.0 nm, Milling current = 0.3 nA, Pattern Type = CleaningCrossSection, Pattern Overlap = 200.0 %, DCM Rescan Interval = 30s), Polishing 1 (Pattern Offset = 150.0 nm, Overtilt = 0°, Depth Correction, 160.0%, High Voltage = 30 kV, Milling current = 0.1 nA, Pattern Type = CleaningCrossSection, Pattern Overlap = 200.0%, DCM Rescan Interval = 30s), Polishing 2 (Pattern Offset = 40.0 nm, Overtilt = 0°, Depth Correction, 160.0%, High Voltage = 30 kV, Milling current = 30 pA, Pattern Type = CleaningCrossSection, Pattern Overlap = 200.0%, DCM Rescan Interval = 30 s).

#### Transmission electron microscopy

Cryo-EM data were collected on a Titan Krios G3 (Thermo Fisher Scientific) operated at 300 keV equipped with a K3 detector and BioQuantum energy filter (Gatan) with 20 eV slit-width. Tilt series were acquired at a pixel size of 2.64Å/pixel at 33000x magnification with tilt range of +/− 44° and 2° increments (46 tilts) using the dose-symmetric scheme. Data was acquired automatically using SerialEM v4.3^32^ with parallel cryo electron tomography (PACE-tomo) scripts^33^ and a 5 µm nominal defocus. For the WT ΦKZ and *imp7 Q4X* dataset, tilt series were collected at 2.17 e-/Å^2^.tilt (Dose = 22.5 e-/pix.sec; 3.22 e-/Å^2^.sec), resulting in a total fluence of 100 e-/Å^2^ (0.67s/tilt). 10 frames were recorded per tilt with 2x binning. All tilt series were processed and reconstructed into tomograms in a single pipeline using AreTomo3 that incorporates MotionCorr and Denoising^34^. Reconstructions were performed with 8x binning.

